# Parieto-frontal Oscillations Show Hand Specific Interactions with Top-Down Movement Plans

**DOI:** 10.1101/2022.05.19.492685

**Authors:** G. Blohm, D.O. Cheyne, J.D. Crawford

## Abstract

To generate a hand-specific reach plan, the brain must integrate hand-specific signals with the desired movement strategy. Although various neurophysiology / imaging studies have investigated hand-target interactions in simple reach-to-target tasks, the whole-brain timing and distribution of this process remain unclear, especially for more complex, instruction-dependent motor strategies. Previously, we showed that a pro/anti-pointing instruction influences magnetoencephalographic (MEG) signals in frontal cortex that then propagate recurrently through parietal cortex (Blohm et al., 2019). Here, we contrasted left versus right hand pointing in the same task to investigate 1) which cortical regions of interest show hand specificity, and 2) which of those areas interact with the instructed motor plan. Eight bilateral areas – the parietooccipital junction (POJ), superior parietooccipital cortex (SPOC), supramarginal gyrus (SMG), middle / anterior interparietal sulcus (mIPS/aIPS), primary somatosensory / motor cortex (S1/M1), and dorsal premotor cortex (PMd) – showed hand-specific changes in beta band power, with four of these (M1, S1, SMG, aIPS) showing robust activation before movement onset. M1, SMG, SPOC, and aIPS showed significant interactions between contralateral hand specificity and the instructed motor plan, but not with bottom-up target signals. Separate hand / motor signals emerged relatively early and lasted through execution, whereas hand-motor interactions only occurred close to movement onset. Taken together with our previous results, these findings show that instruction-dependent motor plans emerge in frontal cortex and interact recurrently with hand-specific parietofrontal signals before movement onset to produce hand-specific motor behaviors.

**Impact Statement:** The brain must generate different motor signals, depending which hand is used. The distribution and timing of hand use / instructed motor plan integration is not understood at the whole-brain level. Using whole-brain MEG recordings we show that different sub-networks involved in action planning code for hand usage (alpha and beta frequencies) and integrating hand use information into a hand-specific motor plan (beta band). The timing of these signals indicates that frontal cortex first creates a general motor plan and then integrates hand-specific frontoparietal information to produce a hand-specific motor plan.

## Introduction

Motor planning is a complex process that encompasses many sensorimotor computations, including sensory processing, target selection, reference frame transformations, and multi-sensory integration (Andersen & Cui, 2009; Crawford et al., 2011). While each of these are important processes, ultimately, a motor plan must be executed using specific effectors (e.g., the eye, hand, or foot). Once an effector system is chosen (i.e., the hand), the brain must still choose *which* hand to use, which motor strategy to employ, and then integrate these signals to produce a hand-specific motor plan. Various studies have investigated hand-target information for visually guided pointing / reaching (see below), but the temporal sequence and frequency-dependence of this process remains unclear at the whole brain level, especially in the presence of ‘top-down’, instruction-dependent motor strategies (Cisek & Kalaska, 2010).

An early step in this process is effector selection (Scharoun et al., 2016), which in the parietofrontal reach system requires hand-specific signals (Filimon et al., 2009). At the sensory input level (somatosensory cortex) there is clear contralateral hand representation, but after that one sees a mix of unilateral and bilateral signals, even down to the level of primary motor cortex (Chang et al., 2008; Donchin et al., 1998; Heming et al., 2019; Matsunami & Hamada, 1981; Tanji et al., 1988; Wiestler et al., 2014). Human neuroimaging studies suggest that parietofrontal cortex is bilaterally activated by unilateral reaches, but with a preference for the contralateral limb (Bernier & Grafton, 2010; Cappadocia et al., 2017; Cavina-Pratesi et al., 2010; Connolly et al., 2003; Filimon et al., 2009; Gallivan, McLean, Valyear, et al., 2011; Gallivan, McLean, Smith, et al., 2011; Gallivan & Wood, 2009; Medendorp et al., 2003; Prado et al., 2005). Likewise, the monkey ‘parietal reach region’ shows some ipsilateral signals (Mooshagian et al., 2018) but is primarily modulated by, and causally related to reaches of the contralateral limb (Chang et al., 2008; Mooshagian et al., 2022). Overall, these findings suggest a progression of ipsilateral and bilateral representations, but the whole-brain distributions and timing of these signals remains unclear.

Further, the presence of hand-specific information (left vs. right hand use, e.g., in somatosensory cortex) does not mean that this has been integrated into a motor plan. Such integration is necessary to activate motor commands for the correct hand, account for the correct initial hand position when calculating the extrinsic hand movement vector, and ultimately activate the correct intrinsic muscle synergies (which will tend to be opposite for opposite arms to produce the same horizontal motion in space) (Gallivan et al., 2013; Ting & McKay, 2007). Most early sensorimotor studies have focused on the feedforward integration of *visual target information* with hand information to compute the reach vector (Khan et al., 2007; Sober & Sabes, 2003). This is thought to occur in parietal cortex (Beurze et al., 2007; Buneo & Andersen, 2006; Chang & Snyder, 2012; Cisek et al., 2003; Gallivan et al., 2013; Hoshi & Tanji, 2000; Medendorp et al., 2005; Vesia & Crawford, 2012). However, it is not known how this occurs in more complex, instruction-dependent or abstract motor strategies (Hawkins & Sergio, 2014; Sayegh et al., 2017).

An example of an instruction-dependent motor strategy is the pro-/anti-reach task where participants are instructed to point toward / away from a visual stimulus (Cappadocia et al., 2017; Connolly et al., 2000; Gail et al., 2009; Gail & Andersen, 2006; Kuang et al., 2016). In our previous magnetoencephalography (MEG) study (Blohm et al., 2019) we found that the pro/anti instruction first influences frontal cortex, in both alpha and beta bands, and then propagates this to more posterior cortical sites. It has been speculated that this might involve ‘mirroring’ the reach goal, which would then require recalculating the reach vector relative to hand-specific signals (Cappadocia et al., 2017; Fernandez-Ruiz et al., 2007; Gail et al., 2009). But again, it is not known how this strategy is integrated with hand-specific information to implement a specific reach command.

In the current study, we recorded MEG signals in the pro/anti pointing paradigm used in our previous study (Blohm et al., 2019) but tested the left and right hand separately. This allowed us to derive both a hand specificity index, and the instruction-dependent motor vector (Blohm et al., 2019). We then performed a region-of-interest analysis based on the areas identified in our previous study (Alikhanian et al., 2013). We hypothesized that although many cortical areas might show hand-dependent modulation (e.g., M1, S1), only those involved in integrating hand information into the motor plan would show an interaction between hand-specificity and extrinsic motor vector coding in their oscillatory activity. Further, we hypothesized that a top-down motor instruction might require recalculating the motor vector, thus requiring a specific progression toward an integrated ‘hand-plan’ motor command. Our results confirm left vs. right hand use specificity in various cortical areas and suggest a specific spatiotemporal progression from independent hand / motor signals to integrated hand-motor coding.

## Methods

We used MEG to obtain brain signals with high spatiotemporal resolution (Baillet, 2017; Niso et al., 2022) that could inform us about hand use and integration of hand information into movement plans. To achieve this, we asked participants to perform a pro-/anti-pointing task in the MEG, using the left and right hands in separate blocks of trials. We then co-registered MEG sensor locations to individual participants’ heads by using an anatomical MRI recording. This allowed us to perform whole-brain source reconstruction of MEG signals and infer the precise oscillatory activity at specific previously uncovered brain regions involved in the task (Alikhanian et al., 2013). We then compared this activity across trial types, left/right targets/movements and the use of left and right hands to see which brain areas differentially synchronize or desynchronize with respect to which hand is used. These procedures are described in detail below. The data set, experimental conditions and most of the analysis pipeline have been described previously (Alikhanian et al., 2013; Blohm et al., 2019).

### Participants

We recruited 10 participants for a pro-/anti-pointing experiment after informed consent, 9 of which performed the task with both hands (7 males, 2 females, 22-45 years old; see statistical analysis section for details about study design and power). Of the participants performing the task with both hands, 8 reported themselves to have a right hand preference and 1 reported themself to have a left hand preference. We screened participants to ensure none had any history of neurological dysfunction, injury or metallic implants, and all (but one with amblyopia) participants had normal or corrected to normal vision. All procedures were approved by the York University and Hospital for Sick Children Ethics Boards.

### Task

Participants performed pro- and anti-pointing movements in 4 separate sets of trials; 2 of those sets were left and right-hand pointing movements respectively with the forearm in the pronated posture. Each set of trials was composed of 100 trials for each of 4 balanced conditions: combinations of target left/right and pro/anti instruction for a total of 400 trials. Figure 1 shows the experimental task. Trials started with a fixation cross, followed by a 200ms combined spatial/task cue presentation (Figure 1A). Cues could appear 5 or 10cm left or right of fixation (we averaged across eccentricities in the analysis) and were either green or red indicating pro or anti-conditions (color-task associations counterbalanced across participants). 1500ms after cue presentation, the fixation cross was dimmed indicating to participants to make a wrist pointing movement towards (pro) or to the mirror opposite location of (anti) the cue.

**Figure 1:**
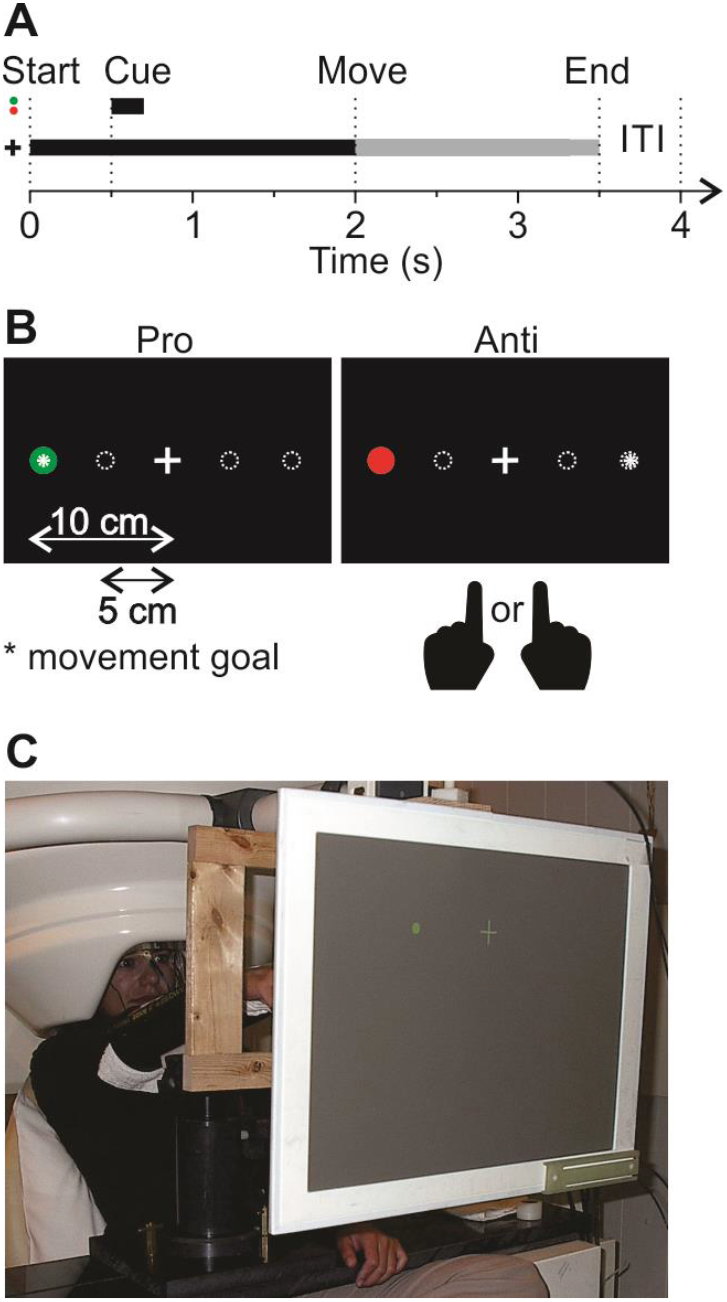
Experimental protocol. **A**. Timeline of experiment. At the beginning of the trial, a central fixation cross appeared. 500ms later, a 200ms cue (red or green) was briefly presented at one of 4 locations, 5 or 10 cm left or right of the fixation cross on the 1-m distant screen. After another 1300ms delay, the fixation cross dimmed to indicate to participants to move and point their index finger to the goal location. Participants had 1500ms to complete this movement, followed by a 500ms inter-trial interval (ITI) during which the fixation cross disappeared. **B**. Spatial setup on screen. Dotted circles are potential cue locations. Exemplary pro- and anti-conditions are shown. In anti-trials the movement goal (asterisk) was at the mirror opposite location of the cue. Pointing movements were performed with the left or right hand in separate blocks of trials. **C**. Picture of the experimental set-up with forearm rest and display screen. The wooden frame held light barriers used for measuring movement direction.

### Set-up

Participants sat upright in the MEG apparatus (151-channel, axial gradiometers, 5 cm baseline, CTF MEG system, VSM Medtech, Coquitlam, Canada, at the Toronto Hospital for Sick Children) in front of a 1-m distant tangential screen with their head in the dewar and their forearm supported to reduce EMG artifacts (see Figure 1C). The MEG was located in a magnetically shielded room (Vacuumschmelze Ak3b). Noise levels were below 10 fT/√Hz above 1.0 Hz. MEG data were online low-pass filtered at 200 Hz using synthetic third-order gradiometer noise cancelation. Bipolar temporal EOG and forearm EMG signals were recorded simultaneously with MEG signals (at 625Hz) to control for fixation and measure wrist movement onset. We used Ag/AgCl solid gel Neuroline (Ambu) electrodes of type 715 12-U/C. Pairs of EMG electrodes were placed over Extensor Carpi Radialis Longior (ECRL), Extensor Communis Digitorum (ECD), Extensor Carpi Ulnaris (ECU), and Supinator Longus (SL) muscles. We also use light barriers that the finger passed when pointing as an additional, independent measure of movement direction (see Fig. 1C).

Visual stimuli were rear-projected (Sanyo PLC-XP51 LCD projector with Navitar model 829MCZ087 zoom lens) at 60Hz onto a translucent screen (Fig. 1C) using Presentation (Neurobehavioural Systems, Inc., Albany, CA, USA) and timing signals were recorded by the MEG hardware through parallel port interfacing. Participants were outfitted with fiducial head localization coils and head position in the MEG was acquired at the beginning and end of each scan. Before or after MEG recordings, we obtained structural (T1-weighted, 3D-SPGR) MRI scans from a 1.5 T Signa Advantage System (GE Medical Systems, Milwaukee, WI), including the fiducial locations for co-registration of MEG signals with brain coordinates (see below; Table 1). For each participant we used the T1-weighted MR data and the BrainSuite software package (Shattuck and Leahy 2002) to derive the inner skull surface.

**Table 1:** Average Talairach coordinates (mm) of functional brain areas. Activation regions of interest were identified using an adaptive clustering approach (Alikhanian et al., 2013) and cross-validated from the literature (indicated by references). We focused on sites corresponding to visual areas V1/2 and V3/3a, SPOC (superior parietal occipital cortex), AG (angular gyrus), POJ (parietal occipital junction), mIPS (medial intra-parietal sulcus), pIPS (posterior intra-parietal sulcus), aIPS (anterior intra-parietal sulcus), SMG (supramarginal gyrus), STS (superior temporal sulcus), S1 (primary somato-sensory cortex), M1 (primary motor cortex), SMA (supplementary motor area), FEF (frontal eye fields), PMd and PMv (dorsal and ventral pre-motor cortex). Note that compared to our previous studies (Alikhanian et al., 2013; Blohm et al., 2019) we updated the names of mIPS, aIPS, SMG and pIPS to bring our nomenclature in line with the most recent literature in this area (Baltaretu et al., 2020; Cappadocia et al., 2018).

| ***Brain area*** | ***Left hemisphere*** | ***Right hemisphere*** | ***References*** |
| --- | --- | --- | --- |
| V1/2 | -8, -91, 0 | 7, -89, 1 | (Martínez et al., 2001) |
| V3/V3a | -21, -85, 16 | 20, -87, 15 | (Martínez et al., 2001; Tootell et al., 1997) |
| SPOC | -9, -71, 37 | 10, -77, 34 | (Vesia et al., 2010) |
| AG | -35, -61, 35 | 32, -70, 35 | (Vesia et al., 2010) |
| POJ | -18, -79, 43 | 16, -79, 43 | (Prado et al., 2005) |
| mIPS | -23, -54, 46 | 27, -55, 49 | (Baltaretu et al., 2020; Bremmer et al., 2001) |
| pIPS | -22, -61, 40 | 23, -62, 40 | (Blangero et al., 2009) |
| aIPS | -37, -40, 44 | 37, -44, 47 | (Baltaretu et al., 2020; Nickel & Seitz, 2005) |
| SMG | -43, -35, 49 | 41, -41, 39 | (Baltaretu et al., 2020; Blangero et al., 2009) |
| STS | -45, -57, 15 | 49, -41, 12 | (Grosbras et al., 2005) |
| S1 | -40, -26, 48 | 39, -26, 40 | (Mayka et al., 2006) |
| M1 | -35, -23, 54 | 37, -23, 52 | (Mayka et al., 2006) |
| SMA | -4, -9, 52 | 3, -7, 49 | (Mayka et al., 2006) |
| PMd | -27, -14, 61 | 21, -14, 61 | (Connolly et al., 2007; Mayka et al., 2006) |
| FEF | -28, -1, 43 | 31, -2, 45 | (Paus, 1996) |
| PMv | -50, 5, 21 | 48, 8, 21 | (Mayka et al., 2006) |

### Analysis

All analyses were done in Matlab (The Mathworks, Inc., Natick, MA, USA). To detect movement onset, we first band-pass filtered EMG data between 15Hz and 200Hz and full-wave rectified it. Then we used an algorithm to automatically detect when EMG signals exceeded 3 standard deviations of baseline activity (measured before target cue onset). The first detection time across all four muscles was taken as movement onset time and visually inspected and manually corrected if necessary (<2% of trials). All data were then aligned to both cue onset (−500 ms to 1,500 ms around cue onset) and movement onset (−1,500 ms to 500 ms around movement onset) and extracted for further analysis. Trials with movement direction errors were discarded from further analysis (3.2% of total trials across participants).

We performed MEG source reconstruction using a scalar (zero-noise gain) minimum-variance beamformer algorithm (D. Cheyne et al., 2007, 2008; Hadjipapas et al., 2005; Vrba & Robinson, 2001) implemented in the Brainwave Matlab toolbox (Jobst et al., 2018) and additional custom code. This inverse method has been shown to achieve high localization accuracy under conditions of low to moderate signal-to-noise (SNR) (Neugebauer et al., 2017; Sekihara et al., 2005). All further analyses were conducted in source space. We focused on previously reported independently identified regions of interest (ROIs) from the same data set (Alikhanian et al., 2013; Blohm et al., 2019); briefly, we used adaptive clustering on peak whole-brain activations in time-averaged raw, non-contrasted data to identify reliable clusters of brain activation and determined area labels that most likely corresponded to the clusters from the literature (see references in Table 1). Note that using the raw, non-contrasted data for determining ROIs was an orthogonal approach to our condition-contrasted analyses, making this a statistically valid approach (Kilner, 2013; Kriegeskorte et al., 2009). We then used the beamformer to extract estimated source time courses of oscillatory activity for each trial at the ROI locations (see Table 1). We used those individual trial data to compute time-frequency responses (TFRs) at those ROIs using standard wavelet transforms. For spatial averaging across participants, individual participants’ source activity was transformed into MNI coordinate space using standard affine transformations (linear and non-linear warping) in SPM 8 and then projected onto a surface mesh of an average brain (PALS-B12 atlas (Van Essen, 2005)) using Caret (Van Essen et al., 2001).

As in our previous study (Blohm et al., 2019) our analysis took advantage of the spatial lateralization of information processing in the brain (e.g. Van Der Werf et al., 2008) to highlight our dependent variables and negate irrelevant variables. Therefore, where applicable, we averaged or subtracted right from left targets, right from left movements, right from left hands use, and/or right from left cortical activation (signal power in a frequency band of interest) for a given brain region. This is in line with what has previously been done in recent neurophysiology (Kuang et al., 2016) and neuroimaging (Blohm et al., 2019; Cappadocia et al., 2017; Gertz & Fiehler, 2015) anti-reach studies. We then used these contrasts to highlight specificity with respect to which hand was used, if sensory processing or movement processing dominated, and whether sensory/motor processing signals were modulated by which hand was used (we call this the hand-sensory/motor code interaction effect).

Since M1 is expected to show strongly lateralized hand effects and motor commands, we used this as a test case to develop our specific analysis pipelines, and then applied these pipelines to our other brain areas (see Results). In the case of hand specific coding, we confirmed that for a given region and hand, the change in oscillatory power was largely independent of target/movement direction (Figure 2A) and pro-anti instruction (Figure 2B), so we averaged across these parameters (Fig 2, right column). Note that this had no influence on hand effects (see results) except double the statistical power of our data. We then subtracted the average activity across all conditions in Left hand from Right Hand to obtain a single hand main effect for each brain area.

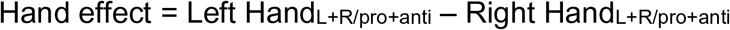

where L/R stands for left/right target location.

**Figure 2:**
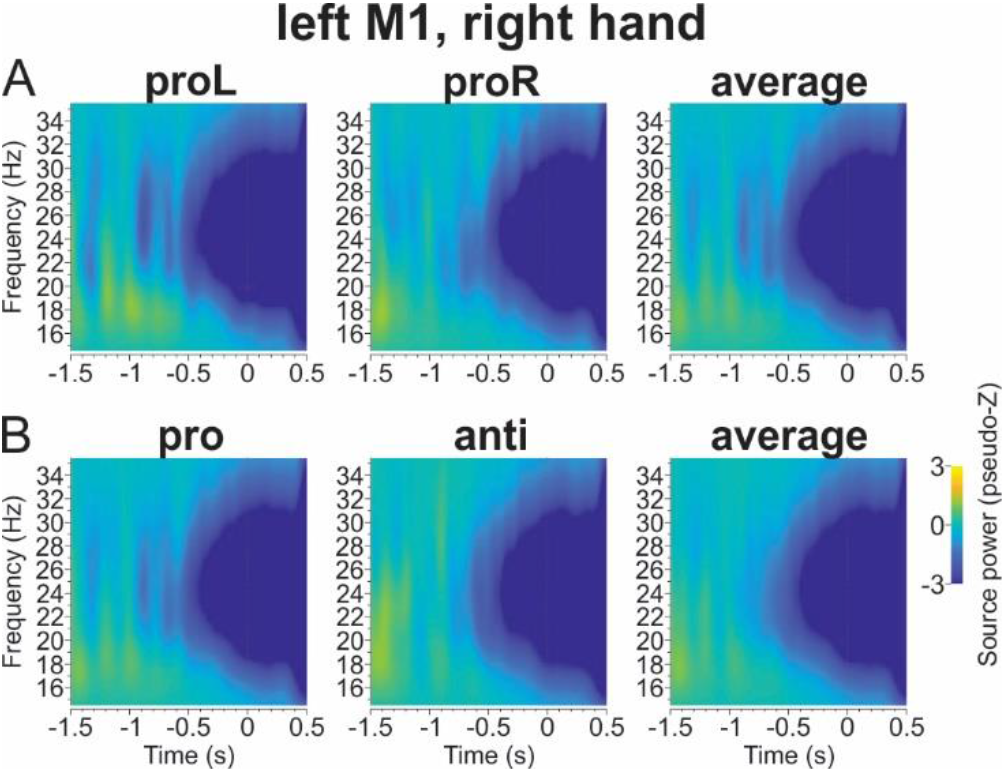
Left/Right and Pro/Anti desynchronization in M1. Time-frequency responses (TFR) for beta band activity (15-35Hz) is shown for right hand conditions in left M1, averaged across all participants. Third column shows the average across A) left and right target/movement directions and B) pro/anti instruction trials. Time zero indicates movement onset.

Finally, in some cases we subtracted left brain from right brain data for each region of interest, to obtain a single bilateral measure of hand effect. The latter parts of this pipeline are summarized in the Results section (Figure 3).

**Figure 3:**
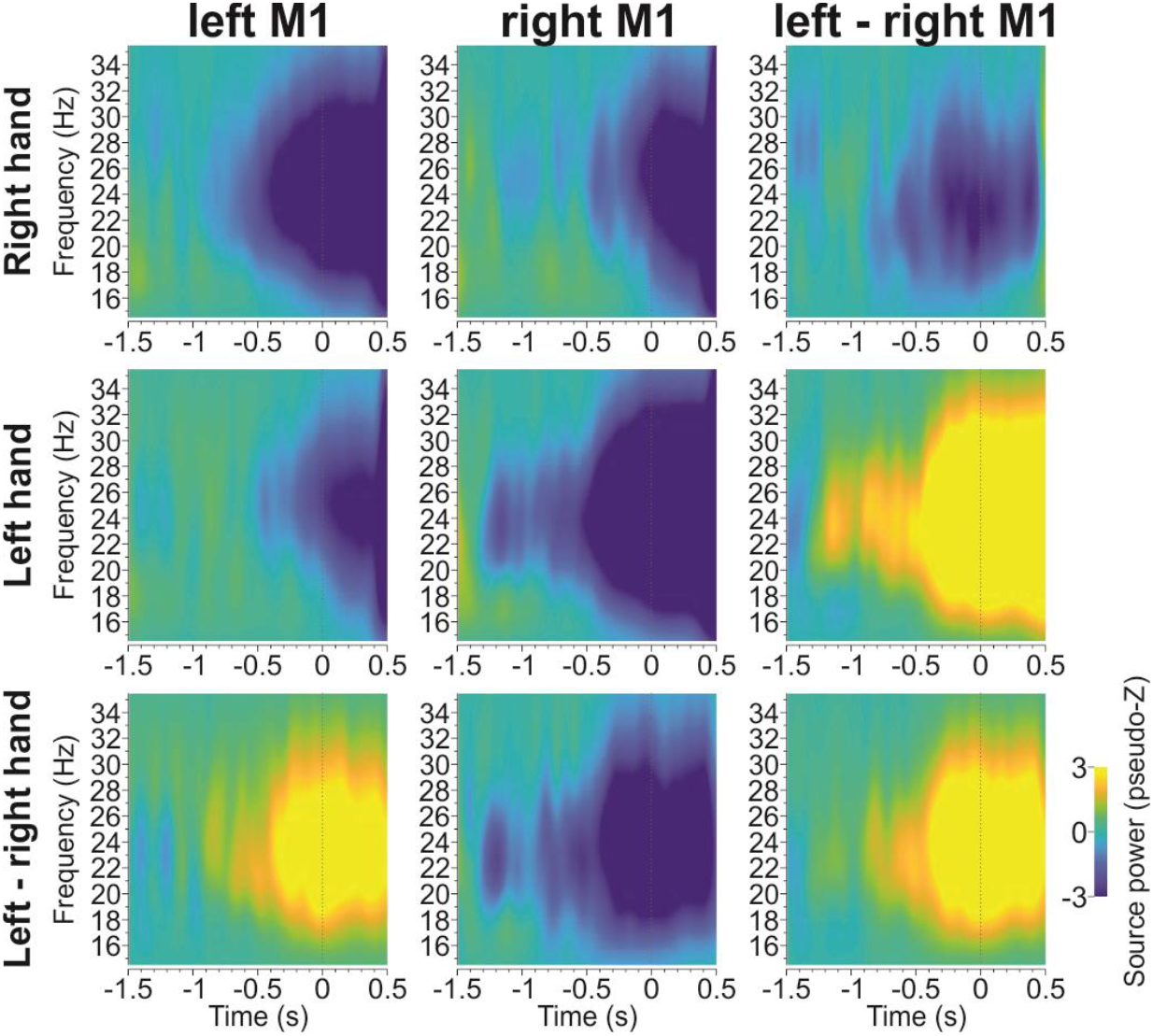
Subtraction logic leading to single bilateral site-specific hand coding index for M1. Each panel shows the oscillatory power change with respect to baseline, averaged across all task conditions (pro/anti, left/right cue) and plotted as a function of time (where zero = movement onset). First column shows activities for left M1, second column shows activity for right M1 and third column shows the differential activity between right and left M1. First row shows right hand activity, second row shows left hand activity, and third row shows the differential activity between right and left hand. The differential activity across sites shows how for a given hand (rows) left and right M1 differentially code for hand information. Conversely, differential activity across hand usage shows how for a given lateralized brain site (columns) the activity differs between left and right hand usage.

To investigate the interaction effect between hand use and motor coding we first computed the spatial motor code for a given hand used, as in our previous study (Blohm et al., 2019):

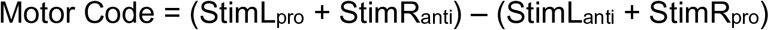

where StimL/StimR correspond to the left and tight sensory cues. Thus, the motor code relies on the fact that the same movement results from a left cue in the pro-condition and the right cue in the anti-condition and vice versa. Note that this procedure is designed to identify a high-level extrinsic spatial code; the motor system must ultimately convert this to hand-specific intrinsic muscle codes (Kakei et al., 2001).

If motor coding dominated an area’s activation pattern, then we expect any trials leading to the same movement to result in similar brain activation; opposite movements should lead to opposite activity patterns, e.g. desynchronization vs resynchronization.

To obtain the interaction between this motor code and hand specificity, we performed the following subtraction:

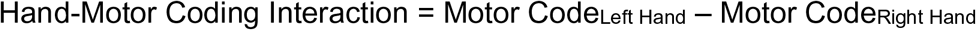

A significant hand-motor coding interaction effect means that the motor code is different depending on which hand is used, indicating integration of hand choice into the movement plan. Conversely, no difference would indicate that the motor code is independent of which hand will be used.

We also computed a sensory code using similar principles (Blohm et al. 2019) highlighting the differential activation of left vs right targets irrespective of movement direction, and tested its interaction with hand specificity:

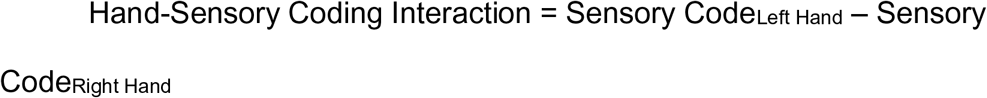

with Sensory Code = (StimL_pro_ + StimL_anti_) – (StimR_pro_ + StimR_anti_)

However, these results were never significant and are not reported below.

### Statistical analysis

Reach studies yield highly robust neuroimaging data and thus tend to employ less participants than perceptual or cognitive studies (Blohm et al. 2019; Cappadocia et al. 2017). In our design, we further offset participant numbers by employing a very high number of trials per participant (Chaumon et al., 2021), e.g. all hand main effect and hand-motor code interaction effect computations relied on the use of ∼800 trials / participant. Statistical significance for individual ROI time series was determined when baseline subtracted source power across participants for a given frequency band was consistently different from zero for at least consecutive 100ms (temporal clustering, (Maris & Oostenveld, 2007)). We used a 2-sided t-test to check for significance (alpha = 0.05). Post-hoc power analysis (G*Power) indicated power > 0.5 (uncorrected for multiple comparisons) for alpha and beta band results in all ROIs, consistent with the standards reported in (Chaumon et al., 2021). Only ROIs that met this criterion are reported below. Note: we did not perform statistical testing for whole-brain averages, as those had been used to identify the ROIs in the first place through adaptive clustering (Alikhanian et al., 2013). Also, adaptive clustering used the raw, non-contrasted, time-averaged whole brain activations and this procedure thus provided independent, orthogonal results to our condition-contrasted analysis here; this prevented double-dipping (Kilner, 2013; Kriegeskorte et al., 2009). Thus, the whole-brain projections are for visualization only.

## Results

### Overview and Predictions

We tested16 bilateral ROIs (see Methods, Table 1), thought to be involved in the sensorimotor aspects of reach planning. Specifically, we investigated visual areas V1 / V2 and V3 / V3a, SPOC (superior parietal occipital cortex), AG (angular gyrus), POJ (parietal occipital junction), pIPS (posterior intra-parietal sulcus), mIPS (medial intra-parietal sulcus), aIPS (anterior intra-parietal sulcus), SMG (supramarginal gyrus), STS (superior temporal sulcus), S1 (primary somato-sensory cortex), M1 (primary motor cortex), SMA (supplementary motor area), FEF (frontal eye fields), PMd and PMv (dorsal and ventral pre-motor cortex). We henceforth use the terms ‘sites’ to refer to the specific coordinates of these ROIs, and ‘areas’ for the surrounding regions.

As noted above, we previously showed that during the pro/anti-reach task, this network of sites / areas first propagates feedforward sensory information (target direction independent of movement direction) from occipital to frontal areas. Sensory coding is present when brain activity patterns for right cue locations are different from left cue locations, but pro and anti conditions do not differ for a given cue location. Next, the instruction-dependent movement plan (movement direction independent of target direction) emerges as the same activity pattern for left cue pro and right cue anti conditions (and vice versa) because they result in the same final leftward (resp. rightward) movement. This instruction-dependent movement plan progressively dominates network activity in a front-to-back progression until movement onset (Blohm et al., 2019). This progression was observed in both the alpha and beta bands, so the same bands were investigated here. For the current study we investigated both sensory and motor code interactions with hand position, but did not find significant hand-sensory interactions, so only hand-motor interactions are reported below.

Specifically, we first describe the main effect of hand use on our sites / areas, and then the interaction of the movement plan (independent of target direction) with hand use to produce an integrated ‘hand-plan’ movement command (Scharoun et al., 2016). To describe hand specificity, we will show contrasts of event-related activity when the left vs the right hand was used while averaging across all stimulus conditions (left/right cue, pro-/anti-trials). We predicted that early hand sensory areas (S1) and late motor areas (M1) would show hand specificity, and asked which intermediate high-level sensorimotor areas would show the same. To investigate the interaction between hand specificity and the top-down motor code we simply subtracted the right-hand motor code from the left hand motor code. We predicted that only late (i.e., after S1) hand specific areas would also show hand-plan motor interactions, including areas that might normally be involved in hand choice for specific tasks and hand-specific conversions from extrinsic to intrinsic muscle coordinates (Kakei et al., 2001). Specific results are described below.

### Hand main effect

Figure 3 shows the later stages of our analysis pipeline (after direction and instruction averaging, see methods) and main result for one example site (M1). M1 is shown here as a site that can be ‘safely’ expected to show contralateral hand dominance, if our method works. Beta band related power changes were averaged across all trials and plotted as a function of time, aligned at the point of movement onset (similar alpha band results will be summarized below). Note that in this first analysis, trials with left/right targets and pro-/anti-pointing instruction were pooled (∼400 trials/hand/participant). The data were first plotted separately for the left / right hand (top two rows) and left / right M1 (left and center columns), corresponding to the four upper-left panels in Fig. 3. Note that lower power (desynchronization, shown as dark blue areas) is associated with *increased* neural activity (Pfurtscheller & Lopes da Silva, 1999). Planning-related power modulations appear to emerge about 1-0.5 seconds before movement onset (zero on the x axis).

To isolate the hand effect and extract a single hand main effect for each site we subtracted right hand trials from left hand trials (bottom row), resulting in what we called the hand main effect. Here, (in the first two panels of the bottom row) yellow signifies more activation for the left hand, and dark blue signifies more activation for the right hand. Finally, we subtracted the left brain from right brain data (right column) to obtain a single measure of bilateral hand specificity for each region (in this case M1). For the top two rows, this subtraction results in the differential activation of left and right M1 separately for the right hand and the left hand. For example, the top row shows that planning to move the right hand leads to stronger desynchronization of left M1 then right M1, as expected. This observation is reversed for planning to move the left hand (center row). For the bottom row, the combined subtraction [Left M1 (left-right hand) – Right M1 (left-right hand)] (Fig. 3, lower right panel), summarizes the overall lateralization of the bilateral structure, where yellow indicates contralateral hand sensitivity. This highlights the expected lateralization of hand coding in M1, with relative left hand modulations in right M1 and right hand modulations in left M1.

We performed the same beta-band analysis on all 16 bilateral pairs of our selected brain sites (Table 1). Of these, we found significant hand-specific activity in 8 bilateral pairs POJ, SMG, S1, mIPS, SPOC, aIPS, M1 and PMd. The data were generally equal and opposite between bilateral pairs (see Supplementary Figure 1), yielding summated power in the final bilateral subtraction (illustrated by the 8 panels in Figure 4). Although each area showed significant hand-specific activation changes prior to movement, they showed area-specific activity patterns. Some sites (M1, S1, IPS, aIPS, PMd) showed relatively well-organized band-time patterns (presumably indicating strong hand preference), whereas others showed intermediate (mIPS) or relative weak patterns (SPOC, POJ). In the latter cases, the onset of hand specificity is not clear from visual inspection of these plots, requiring further quantification (we will return to this point below).

**Figure 4:**
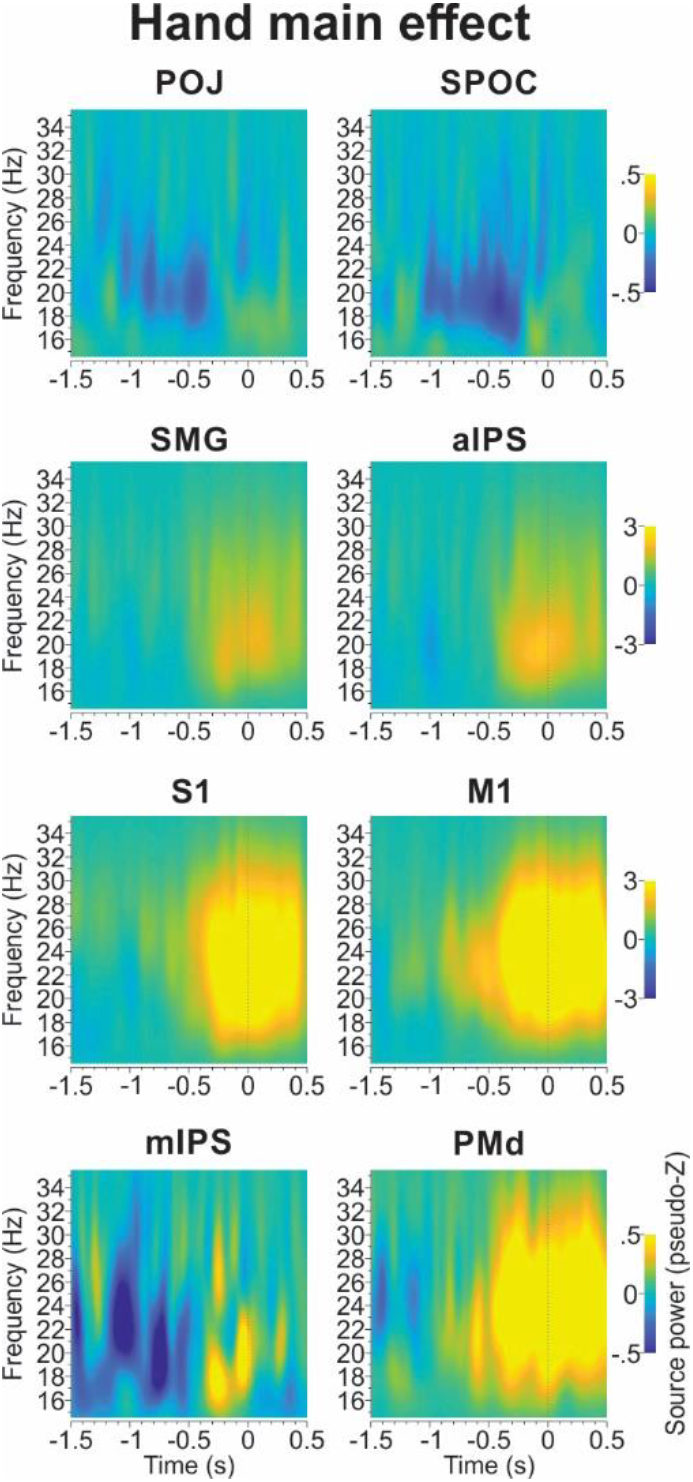
Hand main effect analysis for beta-band activity. Each panel contains temporal plots of activity changes from baseline, averaged across all task conditions. Only shown are the 8 regions that showed significant activation in the overall contrasts between right / left hand and left / right hemispheres, computed in the same way as Figure 3 (bottom right panel).

To visualize these patterns across the entire cortex, we computed average hand-specific beta activity changes from baseline (averaged across conditions) during the last 500ms window preceding movement onset. The result of this analysis is shown in Figure 5 for the left hand (first panel) and right hand, (second panel) separately. (In this case, we removed the left-right hand and left-right brain subtractions, so that whole brain results and their hand lateralization patterns can be viewed.) This shows that prior to movement onset there were widespread changes in oscillatory beta band power across the cortex, with the suggestion of hand lateralization in some sites. To highlight this lateralization, we then subtracted right from left hand average activity patterns (Fig. 5, third panel). This shows strong hand-specific lateralization in dorsolateral parietofrontal cortex, extending slightly more posterior on the left hemisphere (just ahead of our AG / pIPS, right mIPS coordinates) and forward toward bilateral prefrontal cortex (∼ left / right PMv, FEF).

**Figure 5:**
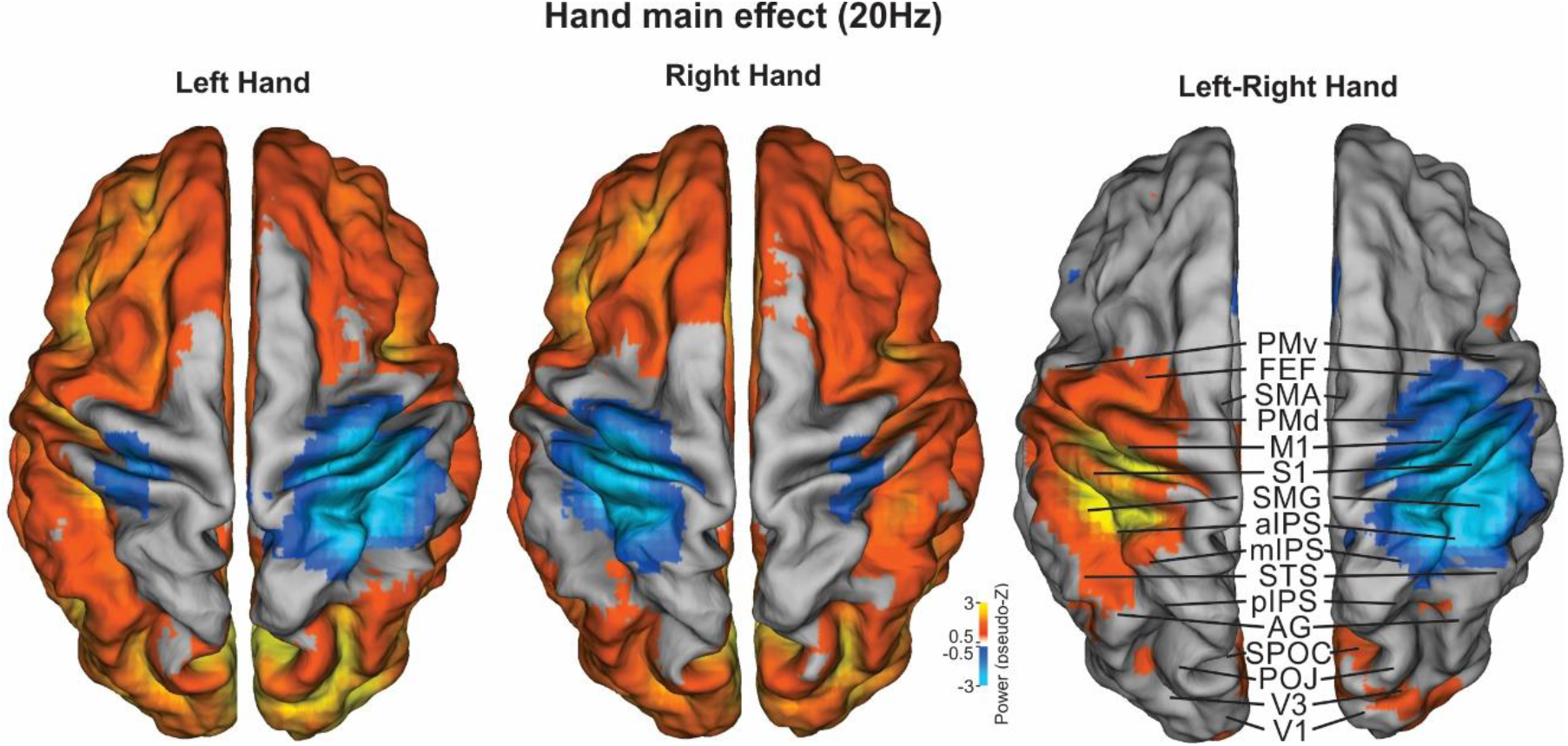
Average movement-aligned activities for each hand and hand main effect (left – right hand subtraction) across cortex. Beta-band activity change compared to pre-task baseline was averaged for the last 500ms prior to movement onset. Subtraction between right and left hand activity (right panel) highlights the hand-specific change in oscillatory power across the cortical surface.

Figure 6 summarizes these observations and extends them both to the alpha band and to the temporal domain. Here, we used the bilateral hemisphere subtraction and then extracted the 10Hz (alpha) and 20Hz (beta) frequencies from the last 500ms before movement onset to generate plots in cortical space (Fig 6, top row). These anatomic plots show peak hand specificity in central regions (mIPS, aIPS, S1, M1) with lesser but still significant activation in surrounding areas (SMG, mIPS, PMd). Overall, beta modulations (right column) were more widespread and pronounced compared to alpha modulations (left column). There were also frequency-dependent regional differences, for example power changes related to hand-specific IPS modulations extended more posterior toward AG in the alpha band compared to the beta band, and beta activation extended further into prefrontal cortex.

**Figure 6.**
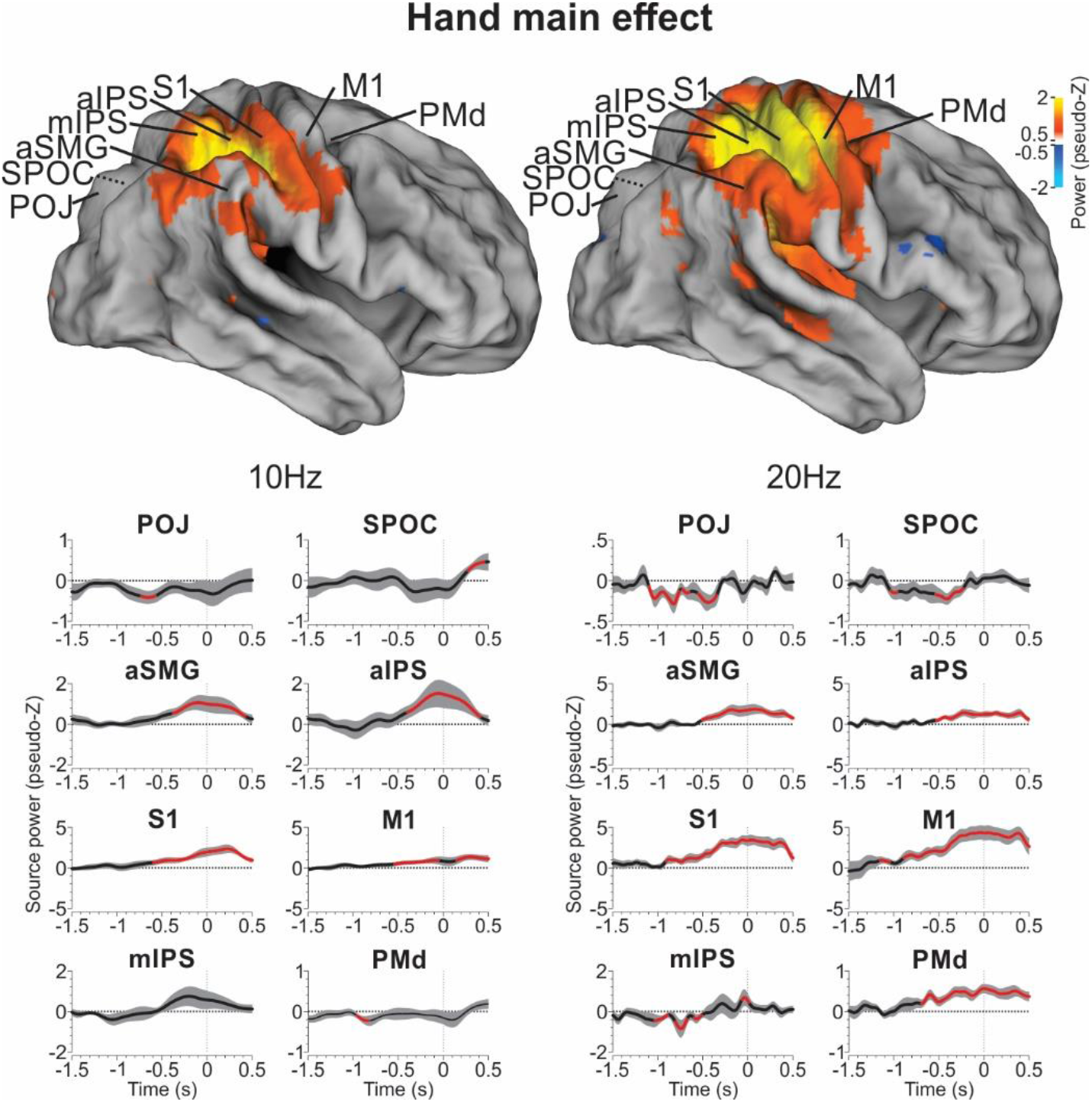
Spatial and temporal specificity of hand main effect in alpha and beta bands. First row shows whole-brain cortical pattern associated with hand specificity averaged during the 500ms prior to movement onset and projected onto the right cortical hemisphere (20Hz data is the same as the third column of Fig. 5). Lower panels show time courses of frequency power (10 Hz left; 20 Hz right) for active brain regions before and around movement onset (time 0ms). Black shows mean signal across participants, grey area is the 95% confidence interval and red indicated significant difference from zero.

The lower rows of Figure 6 show temporal plots of power in the alpha and beta bands for our sites of interest, again aligned on movement onset, where red indicates a significant deviation from equal-hand specificity and a positive deflection indicates contralateral hand specificity. Sites that showed significant contralateral preference typically did so approximately 1-0.5 before movement onset (time zero). SMG, aIPS, S1 and M1 then showed a consistent build-up of significant contralateral hand specificity in both frequency bands during motor planning and execution. PMd showed a similar pattern, but only in the beta band. mIPS showed this pattern in the alpha band, but did not reach significance. Otherwise mIPS and SPOC/POJ activation (which was too small to show up in the anatomic plot) showed oscillations that only transiently reached significance, and in the negative (ispsilateral hand) direction. No significant activations were found in sites V1, V2, V3/3a, AG, pIPS, STS, SMA and PMv.

### Hand-motor coding interaction effect

We previously found that prior to movement onset many parietal and frontal areas displayed coding of the movement plan (see Methods; and Blohm et al., 2019). Specifically, we showed a main effect of motor coding in the same dataset for POJ, SPOC, SMG, aIPS, S1 and M1, but not for mIPS (Blohm et al., 2019). These motor codes were obtained by subtracting data from pro-pointing and anti-pointing trials (pooled in the hand analysis above) such that the directionality of the sensory stimuli cancels whereas the target *and* instruction-dependent motor directions summate (Blohm et al., 2019; Cappadocia et al., 2017). In the previous section of the current paper, we showed that a subset of those areas also show a main effect for coding of hand selectivity (Figures 3-6), but this does not necessarily mean that this information is actually integrated into the motor plan. Here, we tested each of our relevant sites for an interaction effect between the main hand coding effect (described above) and motor coding (obtained in the same was as (Blohm et al., 2019)), i.e., a whether there is a hand-specific motor code.

Fig. 7. shows the steps taken in this analysis and main result, again using M1 as our example site. This figure follows the same steps and logic as Figure 3, except now we are looking at the power of the interaction between the hand effect and the motor code. Similar to the hand main effect, we found roughly opposite patterns of oscillatory power changes when left and right M1 activity were subtracted (top two rows) or when left-right hand activity were subtracted (left two columns). Combining both subtractions (lower right panel) produced a hand-motor plan interaction index for bilateral M1. The significant negative deflection (blue) shown here provides a visual benchmark for what data should look like if a site shows a motor plan specific to the contralateral hand.

**Figure 7:**
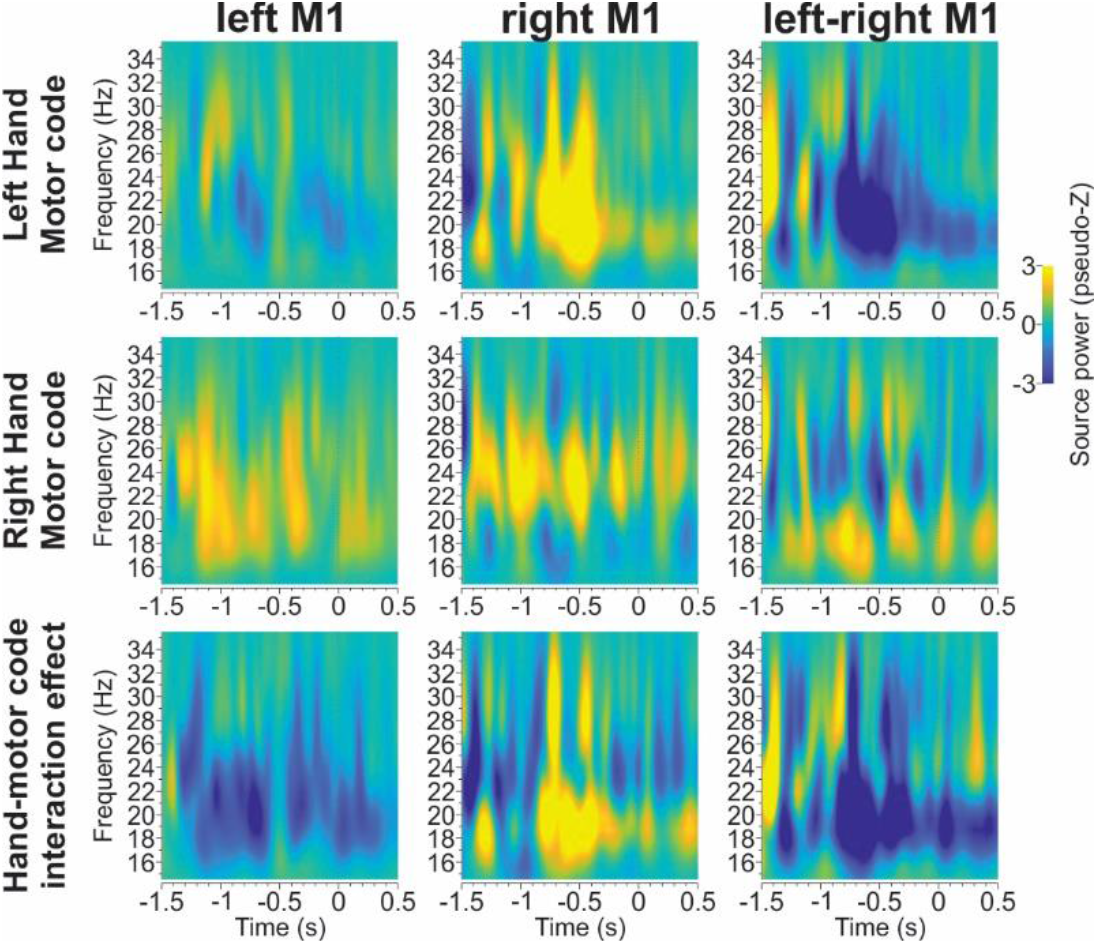
Hand-motor code interaction effect for M1 beta band activity. Time-frequency response plots are shown separately for left hand motor coding (first row), right hand motor coding (second row) and the difference between left and right motor coding (third row) showing the interaction effect for left M1 (first column), right M1 (second column) and the left/right M1 contrast (third column).

Figure 8 shows the result of this beta-band analysis for the 8 bilateral pairs that showed significant hand effects in our previous analysis. Again, blue (desynchronization) corresponds to the expected interaction resulting from both contralateral movement and contralateral hand coding, whereas yellow would denote an opposite interaction: e.g. ipsilateral hand for contralateral motor coding or contralateral hand for ipsilateral motor coding. Note that desynchronization in the interaction effect could also arise from ipsilateral hand and ipsilateral motor coding. Some sites showed a strong interaction in the expected direction (e.g. M1, SMG), others appear to show a mix of positive (blue) and negative (yellow) interactions depending on time and frequency band (SPOC, mIPS, aIPS, S1), and other sites show little or no hand-motor plan interaction effect at all (e.g. POJ, PMd). Of these, only SMG, aIPS and M1 showed clear interaction effects in the 500ms period preceding action, with stronger motor coding for the contralateral hand. SPOC also showed a some interaction, but this was a preference for motor coding in the *ipsilateral* hand. The other sites did not reach significance. (See Supplementary Figure 2 for corresponding lateralized analysis results)

**Figure 8:**
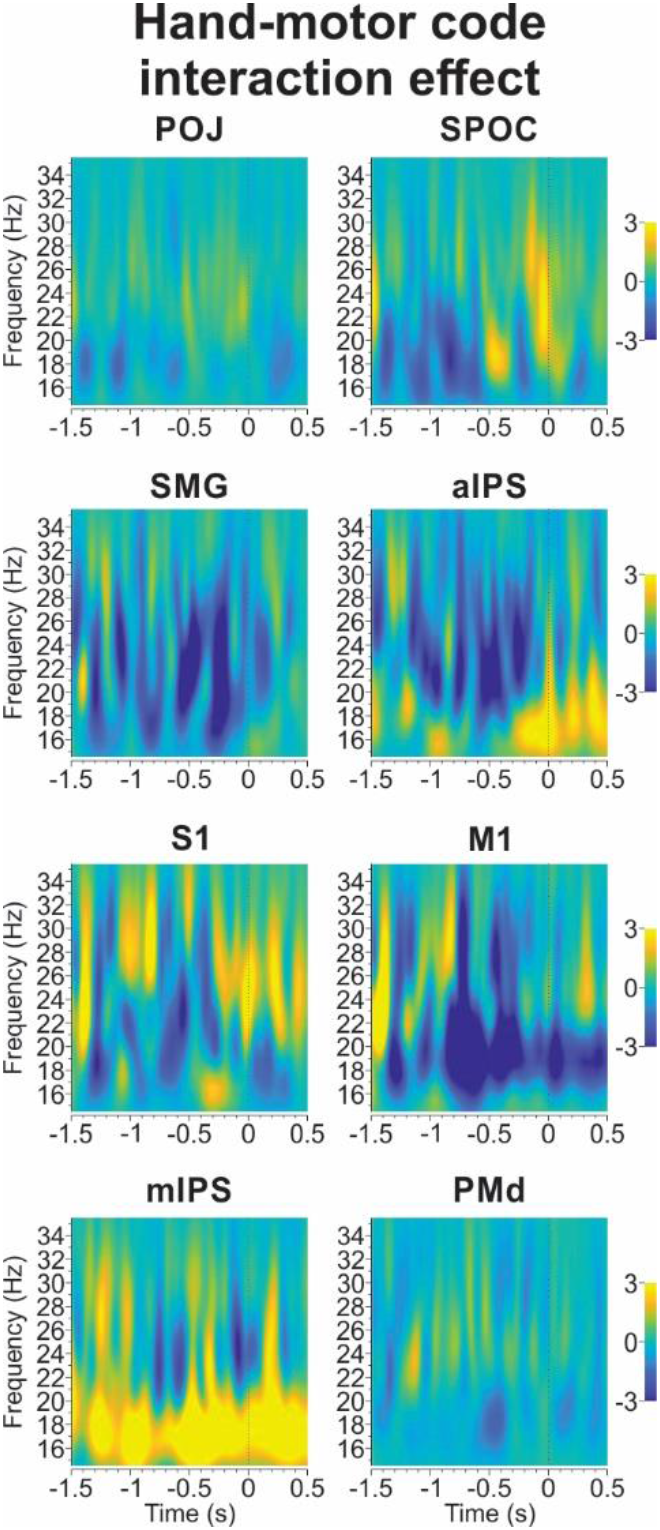
Lateralized hand-motor plan interaction effect analysis for beta-band activity. Same conventions as in Fig. 4 apply.

Fig. 9 shows the functional anatomy and time courses of these interactions in our specific sites, collapsing across hemispheres as described above. The top panels of Fig. 9 show desynchronization related to the interaction effect for 10Hz (left) and 20Hz (right) across the whole brain during the last 500ms time window before movement onset. The beta band shows a broad swath of blue (contralateral hand / motor plan interaction), spanning aIPS, SMG, and M1 and with several other outlying sites. In contrast, in the alpha band, only a small area in posterior parietal cortex showed desynchronization. In short, the hand-motor interaction was primarily observed in the beta band.

**Figure 9:**
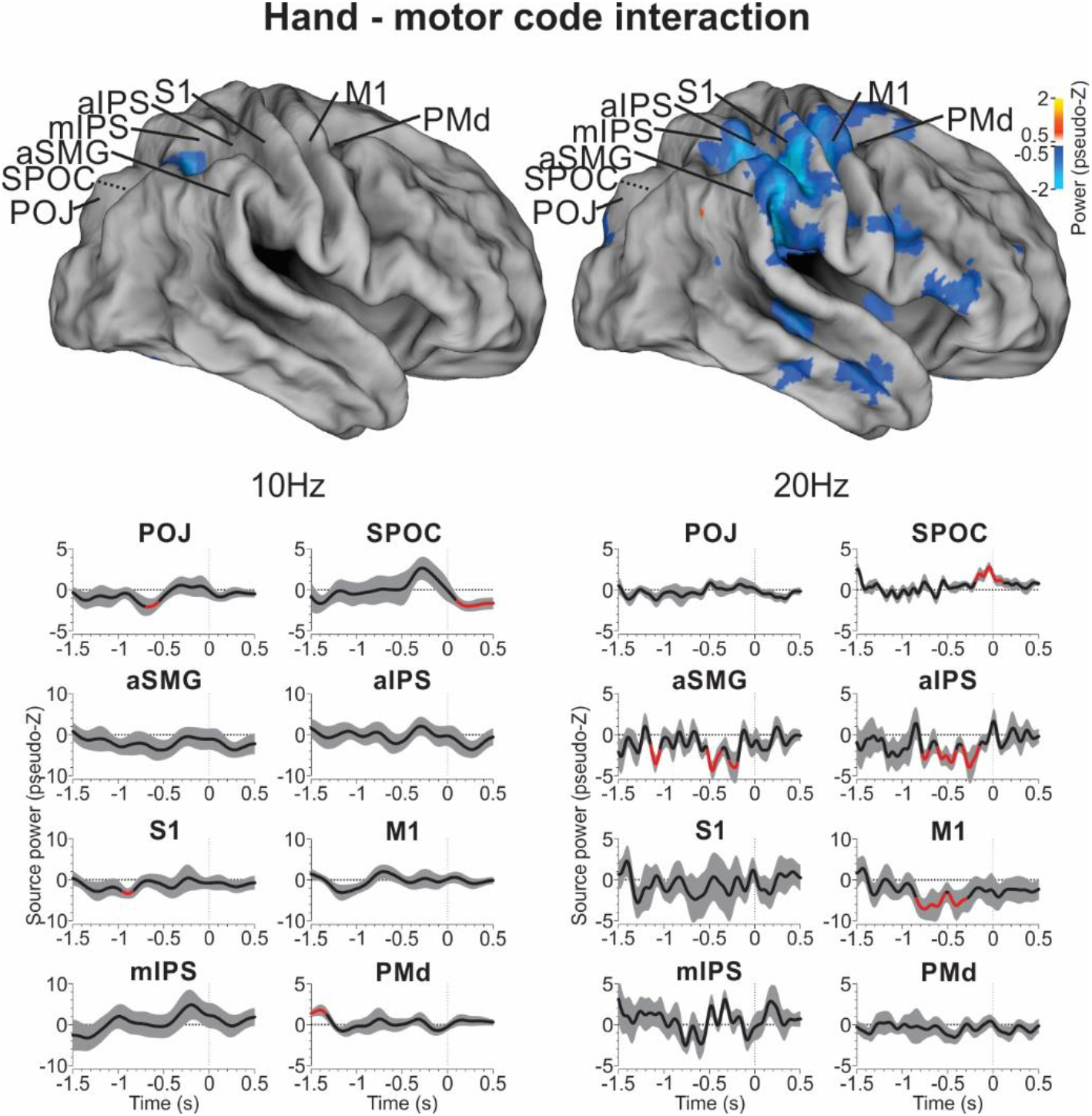
Hand specificity – motor coding interaction. Same conventions as Figure 6 apply.

The lower panels of Figure 9 show a more detailed look at the time course of the interaction effect in both frequency bands, where significant deviations from zero are indicated in red. Consistent with statements above, the alpha band showed very little significant interaction except for a few very brief ‘blips’ before (PMd), during (POJ, S1), or after (SPOC) the planning stage. In the Beta band, aIPS, SMG, and M1 showed significant interactions during the delay period. Interestingly SPOC showed opposite interactions in the alpha and beta bands, and during the movement; this was expected for contralateral motor coding since SPOC showed ipsilateral hand main effects (see Figures 4 and 6).

### Summary: location and timing of hand specificity and motor integration

Figure 10 summarizes the results of both this and our previous study (Blohm et al., 2019). Figure 10A shows the locations of the 16 bilateral sites that we studied, including 13 that showed instruction-dependent motor coding in our previous study (purple outer circles), 8 that showed significant hand-dependences (dark blue middle circles) and 6 that showed significant hand-motor interactions (cyan inner circles) in the current study. It is noteworthy that, while the hand and motor plan networks largely overlapped, some areas (e.g. STS, AG, pIPS, SMA, PMv) only showed movement direction modulations, whereas others (mIPS, S1) only show hand modulation. In general, motor coding was more broadly distributed, whereas hand position modulations and hand-motor interactions (aIPS, SMG, M1) are clustered in superior-medial in parieto-frontal cortex (with the exception of SPOC, with its late ‘opposite’ interaction).

**Figure 10:**
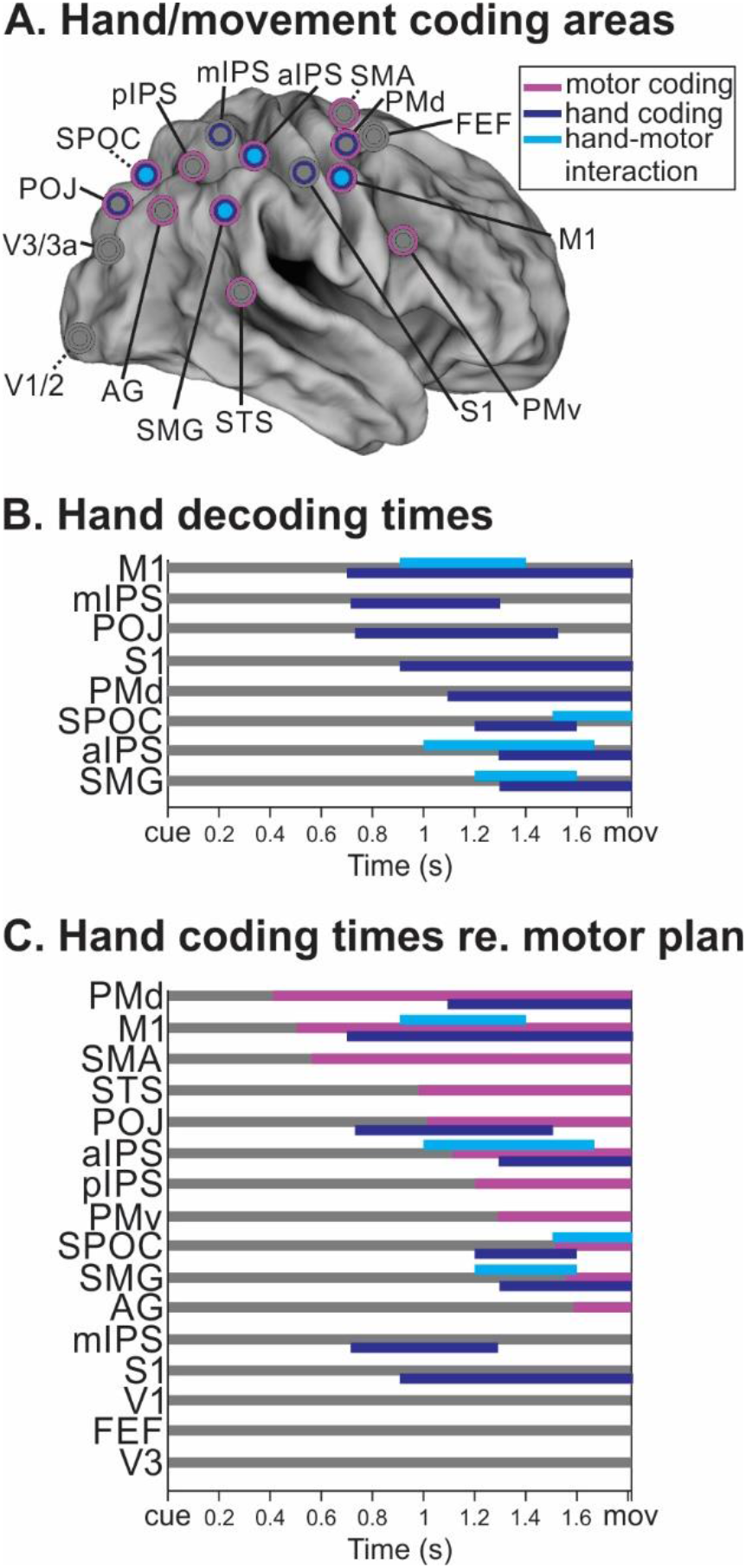
summary of hand specificity and hand-motor code interaction effects. **A**. Colors highlight sites showing significant motor coding (magenta), hand specificity (dark blue) and hand-motor code interactions (cyan). **B**. Times when hand specificity (dark blue) and hand-motor code interactions (cyan) were significant. **C**. Same times as in panel B overlaid with times of motor code significance (adapted from (Blohm et al., 2019)) for all 16 bilateral sites investigated and sorted by motor code onset time.

Figure 10B shows the onset times of hand preference and hand-motor interactions relative to movement onset in the 8 bilateral sites that showed significant hand modulation. Sites M1, mIPS, POJ and S1 showed the earliest hand onsets, but of these only M1 showed an interaction with the motor plan. Sites PMd, aIPS, SPOC, SMG) show later onset, but more interactions with the motor plan (in SPOC, aIPS and SMG). Fig. 10C includes the timing of the top-down motor plan (derived from our pro-/anti-pointing instruction) and all 16 bilateral pairs. As reported previously (Blohm et al., 2019) this plan seems to originate in (pre-)frontal cortex, but is only integrated with hand position in M1 and the more posterior sites described above. Importantly, we see early coding of both the top-down motor plan (in PMd, SMA, M1) and hand signals (M1, S1, POJ, mIPS) but their interactions occurring later, in these and other (aIPS, SPOC, SMG) sites.

## Discussion

We asked how, when and where effector specificity was integrated into instruction-dependent motor commands in the brain using a pro-/anti-pointing task with left and right hand in the MEG. To do this, we performed an ROI analysis on 16 bilateral sites that were previously implicated in instruction-dependent motor planning for this task (Blohm et al., 2019). Our analysis revealed that a sub-set of these areas differentially coded for which hand was used, including robust premotor activity in M1, S1, SMG, and aIPS, with modest but still significant activation in POJ, SPOC, MIPS and PMd. This hand main effect emerged gradually during the pre-motor period and – for some sites – prevailed across the movement (S1, M1 and PMd). Our next analysis found that only SPOC, SMG, aIPS and M1 showed significant interactions of the effector with the movement plan, indicating a role in hand-motor integration (but we did not find significant sensory hand-target direction interactions). As summarized in Figure 10, these interactions occurred within the overlap of two networks that showed early hand / motor signals and later interact close to movement onset.

### Limitations and Caveats

Before considering the physiological implications of these findings, it is worth noting that, like any ROI analysis, our specific sites do not necessarily represent activity in the entire region they are named after, and their locations are best estimates given spatial resolution of the data (MRI/MEG), averaging, and source localization used here. Overall, a reasonable estimate is that these locations are accurate within approximately 5mm, depending on local anatomy (Alikhanian et al., 2013; Chaumon et al., 2021). Given this, one may consider these ROI names and locations as guideposts rather than exactitudes. We will consider more specific aspects of this when we compare our data to the literature below.

A related factor is the relative power and distortions of the MEG signal over gyri versus sulci. In theory, MEG is most sensitive to signals from the walls of sulci for which the cortical surface is orthogonal to the skull surface (Goldenholz et al., 2009; Hillebrand & Barnes, 2002). However, because of current spread, non-spherical skull surface and few cortical surfaces being strictly parallel to the scalp, this is less of a concern in practice (Baillet, 2017; D. O. Cheyne, 2013; Goldenholz et al., 2009; Hillebrand & Barnes, 2002; Koser, 2010). Another caveat is that subtraction methods assume linearity, whereas non-linearities in the MEG signal could either exaggerate or compress differences in activation (Hadjipapas et al., 2005). Next, although MEG has practically unlimited temporal resolution, as in any such study temporal resolution is limited by synchronization with behavioral measures and averaging across participants. Finally, although the number of participants used in this study (performed more than 15 years ago) is low by current standards, our key findings met current standards for power (see Methods). This is likely because sensorimotor tasks yield high and consistent levels of brain activation relative to perception and cognition tasks (D. O. Cheyne, 2013) and because we had many trials for each participant (Chaumon et al., 2021). However, based on these numbers, we cannot draw firm conclusions from negative results. Given these caveats, our findings strongly support our hypotheses and generally agree with the neuroimaging and neurophysiogical literature, as discussed below.

### Instruction-Dependent Motor Codes

It is thought that frontal cortex plays an important role in instruction-dependent and non-standard motor strategies, as opposed to reactive move-to-target strategies (Bonnard et al., 2004; Coe & Munoz, 2017; Hwang et al., 2021). For example, dorsolateral-prefrontal cortex (DLPFC) is thought to be involved in response selection amongst multiple alternatives (Rowe et al., 2000; van Eimeren et al., 2006). Both DLPFC and FEF are involved in preparatory set and task-switching in the oculomotor version of the pro/anti instruction task (Connolly et al., 2000, 2002; DeSouza et al., 2003). Parietal cortex is also influenced by instructions for ‘non-standard transformations’ (Hawkins & Sergio, 2014; Sayegh et al., 2017), specifically causing reversals of motor tuning in primate parietal reach areas (Gail et al., 2009; Kuang et al., 2016). Even occipital cortex appears to be influenced by top-down motor signals from frontal cortex, although those early modulations are thought to involve imagery, attention, gating of sensory inputs, and/or compensation for expected sensory reafference (Gallivan & Culham, 2015; Monaco et al., 2020; Moore & Zirnsak, 2017).

In a previous fMRI study, we provided participants with a pro-/anti-reach instruction following a memory delay, and then used contrasts similar to those used here to isolate visual versus motor directional tuning (Cappadocia et al., 2017). Consistent with the discussion above, we found instruction-dependent modulations of directional tuning throughout occipital-parietal-frontal cortex, with major functional connectivity ‘Hubs’ in superior occipital gyrus (SOG), mIPS, AG, and SPOC, but were unable to determine temporal order. In our subsequent MEG study (Blohm et al., 2019) we provided participants with the pro/anti instruction simultaneously with target appearance and performed a similar directional analysis on the resulting data. As shown here in Figure 10C (magenta lines), this resulted in a sequence of motor recruitment proceeding from frontal toward parietal cortex. These same areas were selected for analysis here, and the motor-vector contrast from that study was used for the hand-motor interaction analysis discussed below.

### Left vs right hand coding

There is evidence for both effector-independent coding (Chang et al., 2008; Donchin et al., 1998; Matsunami & Hamada, 1981; Tanji et al., 1988; Wiestler et al., 2014) and hand specificity throughout the parietofrontal reach system. Even primary motor cortex shows some signals related to the ipsilateral hand (Heming et al., 2019). Human neuroimaging studies show that parietofrontal cortex is bilaterally activated by unilateral reaches, but with a preference for the contralateral limb (Bernier & Grafton, 2010; Cappadocia et al., 2017; Cavina-Pratesi et al., 2010; Connolly et al., 2003; Filimon et al., 2009; Gallivan, McLean, Smith, et al., 2011; Gallivan & Wood, 2009; Medendorp et al., 2003; Prado et al., 2005). Likewise, the monkey ‘parietal reach region’, which spans the medial intraparietal sulcus and area V6A and probably corresponds to mIPS/SPOC (Passarelli et al., 2021) shows some ipsilateral signals (Chang et al., 2008; Merrick et al., 2022) but is primarily modulated by reaches of the contralateral limb (Chang et al., 2008).

In the current study, participants performed blocks of trials with either the left or right hand (and the other hand at rest), so would need to attend to that hand for both sensory purposes, i.e., proprioceptive information about its location (Abedi Khoozani & Blohm, 2018; Burns & Blohm, 2010; Sober & Sabes, 2003) and motor purposes, i.e. to gate motor commands for that hand (Mooshagian et al., 2022; Yttri et al., 2014). Contrasting left and right hand pointing, we found robust premotor specificity in M1, S1, SMG, and aIPS, with modest but still significant activation in POJ, SPOC, mIPS and PMd. Most of these are well known components of the reach system (Gallivan & Culham, 2015; Vesia & Crawford, 2012). Inferior parietal cortex (specifically SMG) is an additional area of interest because it was also activated in our previous fMRI study (Cappadocia et al., 2017), is involved in integrating visuospatial signals for grasp (Baltaretu et al., 2020), and shows anatomic connectivity with temporal, prefrontal, and superior parietal cortex (Vickery et al., 2021).

### Hand-specific movement planning

As noted above, hand-specific information must be integrated into the motor plan, both to account for the correct location and generate motor activation in the correct limb. Functional MRI studies of feedforward hand-target interactions suggest that target and hand-specific information may be integrated as early as parietal cortex, specifically in areas such as mid-posterior intraparietal sulcus (pIPS) (Beurze et al., 2007; Gallivan et al., 2013; Medendorp et al., 2005), although effector-independent signals also may persist within frontal cortex (Wiestler et al., 2014). Brain stimulation studies have suggested that hand-specificity first arises between SPOC and pIPS / AG, the posterior portion of inferior parietal cortex (Vesia et al., 2010). Likewise, electrophysiological studies in monkeys show a progression of hand modulations on visual target signals at the single unit level through parietal and frontal cortex (Chang et al., 2008; Chang & Snyder, 2012; Cisek et al., 2003; Hoshi & Tanji, 2000). We did not find significant hand-target interactions in the current task – perhaps because participants waited to process the pro-anti instruction before developing a movement plan – but cannot dismiss the possibility that these interactions still occurred at a sub-significant level undetectable in our analysis.

Much less is known about the integration of hand-specific signals into top-down motor plans. One potential difference here is there may already be hand-specificity in top-down signals from motor and premotor areas (Scott, 2016). Some of our areas (SMA, STS, PMv, pIPS, AG) showed movement specificity, but not a hand preference. Others (POJ, S1, mIPS, PMd) also code for hand selectivity, but did not show an interaction with the motor code. Hand-motor plan interactions only occurred in a sub-set of areas: SPOC, SMG, aIPS and M1. Hand-target interactions have been reported in monkeys (Passarelli et al., 2021), but these results seem to be at odds with human literature suggesting hand-target interactions in AG and mIPS (Vesia et al., 2010). These differences might be accounted for by the task instruction (i.e. top-down versus bottom-up), specific ROI coordinates (AG is quite large), and/or methodological & spatial resolution differences (MEG vs. fMRI & TMS).

It is noteworthy that our technique only detects extrinsic movement direction (i.e. left vs. right wrist rotation of either the left or right hand). However, the same extrinsic direction of wrist rotation requires opposite (radial/ulnar) deviations in opposite hands. Thus, once this becomes hand-specific, one can deduce which muscles were likely involved. Based on our EMG recordings, this would require specific intrinsic synergies of activation in the Extensor Carpi Radialis Longior, Extensor Communis Digitorum, Extensor Carpi Ulnaris, and Supinator Longus (SL) muscles, as well as other forearm muscles that we did not record from. This switch from extrinsic to intrinsic muscle coding is thought to only begin at the level of M1 (Kakei et al., 2001). This could signify that our interaction results in M1 were related to this extrinsic-intrinsic conversion, whereas SPOC, SMG, and aIPS are likely related to higher level aspects of hand-motor integration. For example, in our task we chose the hand for our participants, but in real world circumstances these areas might be involved in hand choice for specific tasks (Scharoun et al., 2016).

### Timing

Thus far we have discussed the spatial distribution of signals in our regions of interest. For this purpose, MEG might have a lower spatial resolution than fMRI, but an important advantage of MEG is its higher temporal resolution. The relative activation of various signals in our regions of interest (summarized in Figure 10) provides clues to the order of processing in our task. For example, it could be that in the anti-reach task the brain waits until the final motor stage to ‘flip’ the desired reach vector. Alternatively, frontal cortex could ‘flip’ a high-level goal (opposite the stimulus) and then feed this back to hand-specific areas to recalculate the desired motor vector (Cappadocia et al., 2017; Fernandez-Ruiz et al., 2007; Kuang et al., 2016).

The timing of events in our data appears to be consistent with the latter possibility. First, besides visual signals (not shown here) one of the earliest signals that we observed was the instruction-dependent motor signal in frontal cortex, which then appears to feedback recurrently to parietal cortex. The next signal to emerge was hand-specificity, suggesting a role in motor preparation. Notably, both these signals (hand and motor) were sustained through the action phase, suggesting they play roles in both planning and execution (Heming et al. 2019). In contrast, the hand-motor interaction occurred closer to movement onset, suggesting a more specific role in planning (such as calculation of the motor vector). Overall, these results are consistent with the notion that the pro / anti instruction does not just influence the final motor output vector but causes updating and integration of hand-motor signals throughout parietofrontal cortex.

An interesting aspect of the timing of hand-specific motor commands (interaction effects) is that they first show up in M1. Following M1, hand-specific motor commands later also appear in more posterior (parietal) areas. This succession of timing in the coding the motor plan aligns with our previous finding (Blohm et al., 2019) that sensory signals first undergo a feed-forward transformation along the posterior-anterior axis, with motor coding first appearing in M1 and then gradually appearing in more posterior areas again. We have suggested that M1 updates the motor intention in more posterior regions once it has been computed after the initial feed-forward transformation. Here, we show that early hand coding in M1 also leads to an early interaction effect. This hand-specific motor code then gradually appears in more posterior areas (SPOC, aIPS, SMG). Overall, this timing suggests that M1 is the main driver in establishing a hand-specific motor code.

### Frequency specificity

Another advantage of MEG is its capacity to isolate frequency-specific effects. This is interesting because different frequencies are associated with different processes in the brain, e.g. alpha band modulations indicate sensory processing (Buchholz et al., 2014; Jensen et al., 2002; Klimesch, 2012; Palva & Palva, 2007), and beta band modulations accompany motor planning and control (Buchholz et al., 2014; D. O. Cheyne, 2013; Isabella et al., 2015; Kilavik et al., 2013; Lopes da Silva, 2013; Neuper et al., 2006; Neuper & Pfurtscheller, 2001; Spitzer & Haegens, 2017; Van Der Werf et al., 2009). Consistent with sensory coding of hand use, we found a main hand effect in alpha band. We also found a main hand effect in beta band, which makes sense given this was a movement planning task. At the same time, we only observed a significant interaction effect in the beta band, suggesting that this interaction effect reflects the actual motor plan specific to the hand used. At the same time, our previous study (Blohm et al., 2019) showed that an abstract (not hand specific) motor intention was decodable from both alpha and beta bands. While we cannot interpret negative results (i.e. absence of alpha interaction effects), our results are certainly consistent with a sensory-to-motor transformation using sensory hand and target information and transforming it into a hand-specific motor plan.

### Correlation vs. Causality

The ultimate challenge for this area of research is to reconcile the evidence for hand preference for bilateral hand representation in the reach system, even down to the level of primary motor cortex (Ames & Churchland, 2019; Bundy & Leuthardt, 2019; Heming et al., 2019). An important distinction here is between correlation and causality: techniques such as unit recording, fMRI and MEG show the presence of signals, but do not necessarily imply a causal relation to the movement. A growing consensus is that, despite bilateral representation, contralateral causality emerges as early as the parietal reach region (Mooshagian et al., 2022; Yttri et al., 2014). Ipsilateral signals might play other roles such as bilateral coordination (Le et al., 2017; Le & Niemeier, 2014), but could be filtered out for contralateral control through parcellation of signals (Ames & Churchland, 2019; Bundy & Leuthardt, 2019; Heming et al., 2019).

## Conclusion

Together with our previous study (Blohm et al. 2019), these results demonstrate a specific whole-brain topography for hand-specific signals and instruction-dependent motor signals in parietofrontal cortex, in both the Alpha and Beta bands. In contrast, hand-motor interactions primarily occurred in the Beta bands, and in a smaller sub-set of frontoparietal regions. The timing of these events suggests that in an instruction-dependent pro/anti pointing task, the motor strategy (same or opposite to visual stimulus) is determined in frontal cortex and then propagates backwards to parietal cortex, with premotor hand-motor interactions occurring later before movement onset. Generalizing from these results, we suggest that top-down, instruction-dependent and/or abstract motor strategies show a different sequence and topography than bottom-up hand-target interactions in the feedforward occipital-parietal-frontal path.

## Acknowledgments

Experiments were supported by a Canadian Institutes for Health Research Grant held by JDC. GB was supported by a Marie Curie Fellowship (EU) during the experiments and by NSERC (Canada) thereafter. During these studies, JDC was supported by a Canada Research Chair. The authors would like to thank Andreea Bostan, William Gaetz, Herbert Goltz and Sonja Bells for technical assistance during data collection.

## Supplementary Figures

**Supplementary Figure 1:**
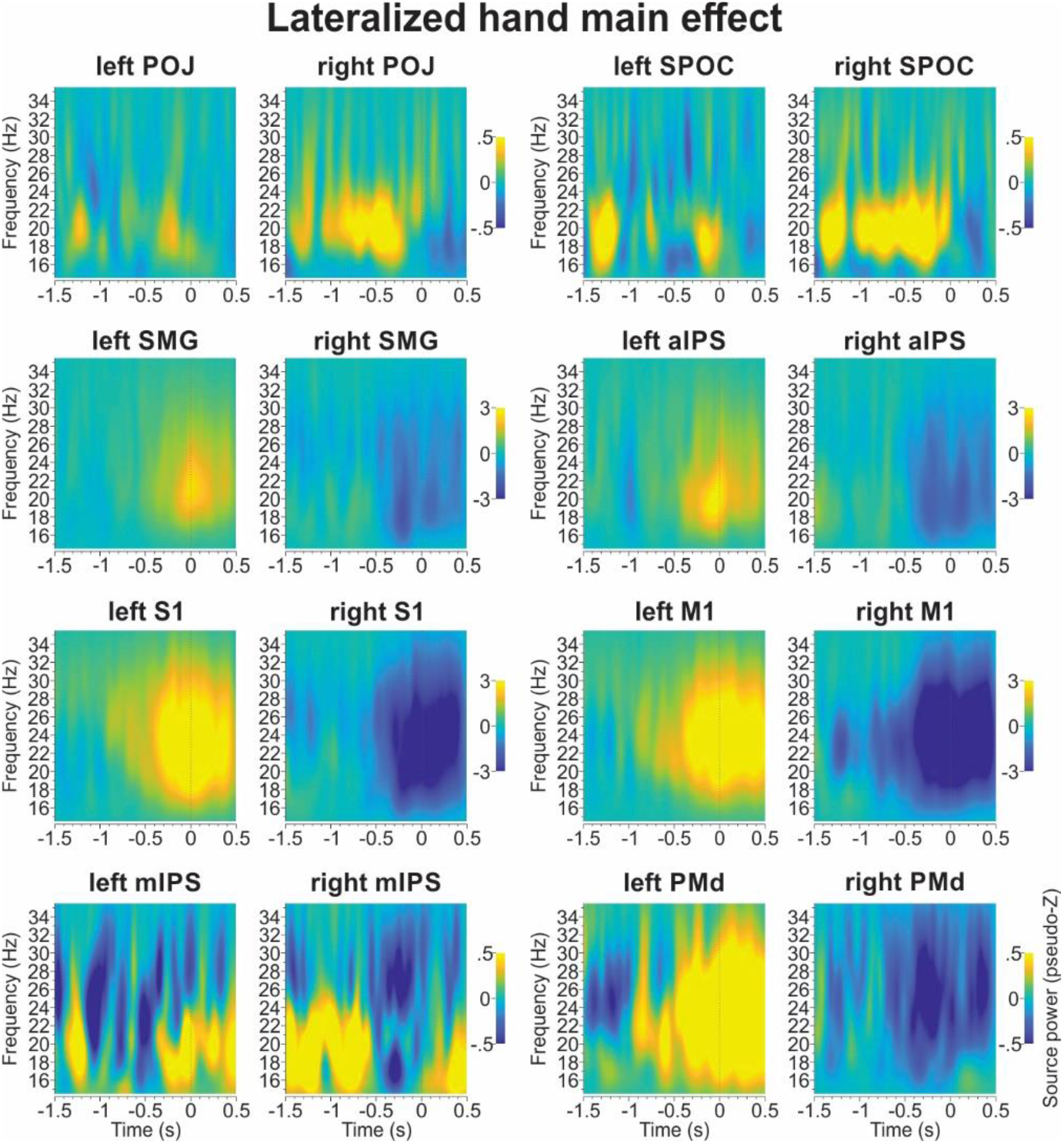
Main hand effect (beta band) separately for the corresponding left and right cortical areas. (same conventions as Figure 4)

**Supplementary Figure 2:**
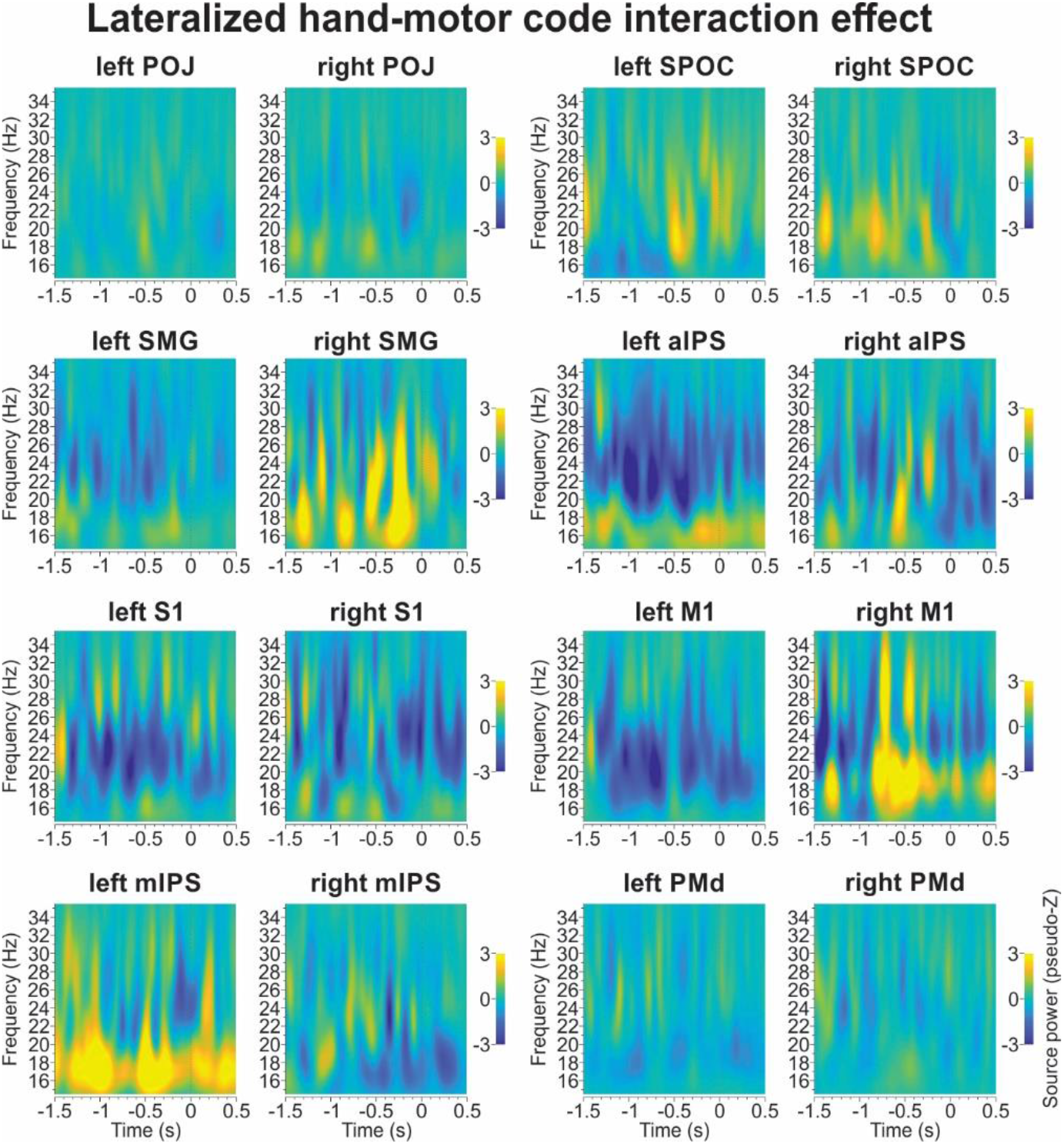
Hand-motor code interaction effect (beta band) separately for the corresponding left and right cortical areas. (same conventions as Figure 8)

